# The Role of Ion Transition from the Pore Cavity to the Selectivity Filter in the Mechanism of Selectivity and Rectification in the K_v_1.2 Potassium Channel: Transfer of Ion Solvation from Cavity Water to the Protein and Water of the Selectivity Filter

**DOI:** 10.1101/2020.03.16.994194

**Authors:** Alisher M Kariev, Michael E Green

## Abstract

Potassium channels generally have a selectivity filter that includes the sequence threonine-threonine-valine-glycine-tyrosine-glycine (TTVGYG). The last five amino acids are conserved over practically the entire range of evolution, so the sequence obviously is necessary to the function of the channel. Here we show by quantum calculations on the upper part of the channel “cavity” (aqueous compartment between the gate and selectivity filter) and lower part of the selectivity filter (SF) how the channel with two sets of four threonines (the channel is fourfold symmetric) effects rectification and selectivity. The threonines are at the location in which the ion transfers from the cavity into the SF; in this calculation they play a key role in selectivity. The channel is also a rectifier. The wild type channel with K^+^ and three other cases are considered: 1) the upper set of four threonines is replaced by serines. 2) and 3) Related computations with the Na^+^ and NH_4_^+^ ions help to clarify the important factors in moving the ion from the cavity to the SF. In particular, one set of angles (not bond angles, **O**(T373–C=O) – **O**(T374–OH) – **H**(T374–OH)) flips a hydrogen into and out of the ion path, allowing the K^+^ to go forward but not back. This is essentially a ratchet and pawl mechanism, with the ratchet driven by electrostatics. This also allows a clear path forward for K^+^ but not for Na^+^ or NH_4_^+^, nor for K^+^ in a T→S mutant. Atomic charges in the lowest positions in the SF are the driving force moving the ion forward, but the O - O - H angle just specified is key to making the “knock-on” mechanism move the ions forward only, using the ratchet with the pawl formed by the hydrogen in the bonds that flip. A water interacts with threonine hydroxyls to allow ion passage, and another water moves together with the K^+^.

## INTRODUCTION

Potassium channels are found in all cells, and in particular are required as part of excitable tissue, including nerve and heart. They have been extensively studied both for their intrinsic importance and because of the diseases that result when the channels do not function normally. There are several properties that practically all channels have, and are certainly important in the voltage gated potassium channels we consider here: gating, selectivity, conductivity, and inactivation. Selectivity is a function of a selectivity filter in the channel pore. The well-known sequence threonine-threonine-valine-glycine-tyrosine-glycine (TTVGYG) in potassium selectivity filters (SF) is found in essentially all potassium channels, certainly voltage gated channels, plus virus (1) and bacterial channels(2, 3) that are not voltage gated. Save for occasional minor mutations of the valine, and sometimes the first threonine, this has been conserved through half a billion years, and probably more, of evolution. Nature having solved the problem of K^+^ selectivity, it has saved the solution.

The first step controlled by the SF is transfer of the ion from the channel cavity which is in the center of the channel pore through which the ion must travel, where it is generally hydrated, to the SF where it must be cosolvated by the amino acids, starting with threonine (we follow customary usage, and call the intracellular end of the SF the lower end). The function of both threonines is generally assumed to have to do with the co-solvation of the ion by some combination of the hydroxyl group and backbone carbonyl groups, but the atomic level details are of critical importance. The problem has been considered by a number of groups, but still is not solved. The reason for the conservation of the threonine, which is rarely substituted by serine, for example, suggests the importance of the geometry and location of the hydroxyl, and requires determination of the exact structure. The wild type (WT) X-ray structure not only of KcsA, a bacterial channel with a small cavity in its pore, but of the eukaryotic, and larger, voltage gated channel K_v_1.2, is known from the work of MacKinnon and coworkers (4, 5). A number of cases have been calculated for the KcsA bacterial channel. We have done such a study for that channel some time ago (6, 7). Many studies of this channel have also been carried out by other workers. Ban and coworkers(8) have presented the most relevant example, in which they attempted to solve two problems: transfer from the hydrated cavity to the lowest section of the SF, and the exit of the ion at the other end, using quantum mechanical (QM) calculations similar to those we use). Their model of the filter consisted of a section at the cavity to filter boundary, but without any of the cavity or its waters, and a section at the extracellular end, with a cylinder without atoms between. They assumed that serine and threonine should be essentially equivalent, and did not compute the actual cavity+lower section structure, but calculated energy assuming full hydration of the ion in the cavity. They did calculate free energy at room temperature for both Na^+^ and K^+^, but based on their hydration assumptions. We obtained somewhat different results in our earlier work (6, 7) as well as in this calculation. Varma et al (9), find the basis for selectivity still not solved a decade ago. Since then, a number of papers that present a variety of calculations and experiments (10-16) have made attempts to consider the problem; this selection is far from exhaustive, as the literature is voluminous, and the approaches varied. Several authors approach the problem from a thermodynamic point of view, some from a structural point of view. The principal experimental techniques include X-ray diffraction and fluorescence resonance energy transfer (FRET). However, a recent solid state NMR study used ^15^NH_4_^+^ as a proxy for K^+^ to determine ion binding density (17) in the SF, to obtain ion binding dissociation constants, with pH dependent occupancy. However, results presented here suggest that NH_4_^+^ is not a legitimate proxy for K^+^; it behaves more like a charged water molecule. A single molecule FRET study finds the SF switching from constrained to dilated conformation, with the constrained conformation, present with K^+^, essential for K^+^ selectivity. Langan and coworkers (18) used anomalous X-ray diffraction to find an average occupancy of 0.25 per site, which they interpreted as meaning that all four sites are fully occupied; as they noted themselves, this contradicts data showing cotransport of water (19, 20). Their claim was that this supports the direct “knock-on” model in which an ion pushes the ion ahead of it forward. More recently, Langan et al used neutron diffraction to clarify the ion to water ratio in the SF (21). Tilegenova et al (22) used two mutants of the KcsA filter to support the ion-water-ion-water sequence in the filter. Kratochvil and coworkers used 2D IR, with MD simulations, in a semisynthetic KcsA channel, to conclude that water separated the ions in the SF(23). However, Strong et al(24)(24) now show that the Kratochvil et al result fails to settle the matter. Weingarth et al combined solid state NMR with molecular dynamics (MD) simulations on the KcsA channel, and found buried ordered water molecules, primarily in the upper part of the selectivity filter, and outside the pore (25). This is a section of the channel we do not consider in our calculations. However, they also found some ordered water, or possibly weakly ordered water, that they attributed to connections of the SF to the remainder of the pore, and these findings are consistent with our calculations, although they cannot be taken as direct confirmation.

There are a large number of simulations and other calculations as well; we consider quantum calculations (QM) first. Cifuentes and Semiao (26) proposed the existence of two paths through the filter, using a time-dependent external field, producing resonances and interference. Summhammer and coworkers (27) carry this one step further in considering the delocalization of the ions. They state that the deBroglie wavelength is not much smaller than the Coulomb potential periodicity. For K^+^ we find this unlikely, although for H^+^ with a deBroglie wavelength λ_D_≈ 0.3 Å, assuming momentum at room temperature equal to h/√(2mk_B_T) (h= Planck’s constant, m=ionic mass, k_B_=Boltzmann constant, T=temperature). However, for the much heavier, and more relevant, K^+^ ion, λ_D_ is about 6 times smaller, and thus should be out of range of the periodicity of the potential. Summhammer et al argue that transient coherence among ions is required to explain the rate of transport of K^+^ ions. Vaziri and Plenio (28) also argue that quantum coherence is important for the SF, and suggest experiments to test this hypothesis. De, March et al (29) extend this argument by including Coulomb repulsion among ions. De et al(30) carried out a QM/MM (MM = molecular mechanics) calculation on the KcsA SF, starting with the lowest threonine, T75 in the 1k4c numbering. They do not include the cavity at all, and the discussion of the role of water is limited. They conclude that each of the amino acids in the TVGYG sequence (the preceding T is not as conserved) has a role to play, with Y78 having key importance in selectivity. Pichierri (31) has calculated the dipole of the KcsA channel and found that the full length KcsA dipole is 403 D (D=Debye), from which he concludes that the channel dipole has a role to play in transport, something no other author has emphasized. IIlingworth and Domene, and Illingworth et al (32, 33) have reviewed results on the effects of polarizability. Pless and coworkers (34) emphasized hydrogen bonds, with connection to inactivation. Miloshevsky and Jordan (35) carried out an energy minimization study, also with relevance to inactivation. Bucher and coworkers (36) studied the effects of coordination number, using a QM/MM method. One paper (Cifuentes and Semiao (37)) actually looked at quantum interference of the ions in the SF, a very different matter than a standard QM calculation, which optimizes structure by finding an energy minimum, although it seems unlikely that such large ions as K^+^ would show interference effects if separated by as much as 5 Å (see the comment on de Broglie wavelength above).

Several authors have considered the role of entropy in selectivity (14), (38-42). Generally, this is attributed to the conformational shrinking or expanding of the filter, as the amino acids make more or less room for the ion. When the ion is constrained the entropy is lower. However, not all studies have found this type of constriction. While the experimental evidence does not entirely rule it out, there does not seem to be very strong experimental support for this type of constriction either (see also Weingarth et al (25) for example for a possible role for ordered water in producing these effects; this said, they allow for alternating conformations of the SF). These authors find several ways in which the constriction of the SF presents entropic barriers to the ion in the SF. They do not agree in detail, but the general conclusion is that constriction lowers entropy, and this can lead to a barrier to the advancement of the ion. Dixit and coworkers (43) considered coordination of the ion, also with consequences for the entropy. QM approaches give accurate optimizations of a section of the protein, but this may not be a section sufficient to provide a good representation of the actual performance of the channel. It also ignores entropy.

The most appropriate form of computation, QM as against MD, is still controversial. Questions have been raised about QM, for example by Roux and others, concerning local minima, the contribution of the surroundings being small but not negligible (44, 45); thermal averaging gives more accurate results, and the local contribution to the energy may, in principle, be important for the free energy(43). In this work, the energy results suggest that local minima are not a problem (further discussion on this point below), surroundings are subtracted in taking differences, but we still do not have results at a finite temperature. The most obvious disadvantage of QM is that the calculation gives the structure at 0 K. It therefore omits thermal averaging and entropic effects. This means that the QM approach, when used for comparison between different ions or mutations, implicitly assumes that the contribution of vibrations of the protein is either neglected, or assumed to be very similar in the cases compared so that it subtracts out in the differences used for data here. It is small compared to the main energy terms, but large enough that it can make a difference in some cases, as has been pointed out, for example, by Yu and Roux (46). Here, the number of degrees of freedom affected by the differences in structure is much smaller than the total, so the differences should be small. However, differences of a few kJ cannot be ruled out, and this may be greater than k_B_T (≈2.5 kJ at 300 K). Most of our conclusions are based on structural calculations; energy calculations are consistent with these, but the main conclusions do not hang on them.

The most common form of calculation on the filter, as on the rest of the channel, is MD (44, 46-48). Various results, and models, have resulted from these calculations, depending on the starting conformation, on the choice of force fields, and other parameters. Several of the experimental papers have been supplemented by MD calculations. Bucher and Rothlisberger commented on MD from a quantum point of view. Because the results are so variable, interpretation is not obvious. Bucher et al(36) showed, using a QM/MM calculation, that the correct coordination numbers of ions in the channel are not the same as those derived from MD calculations. We will not consider MD calculations further, as the results do not seem consistent, the force fields may not be accurate, and polarization along with electron shifts that alter local charges as the ions move is usually neglected.

We may briefly compare the K^+^ selective channels to the non-selective NaK channel, which allows all monovalent, and, apparently, some divalent, ions to pass. There is a large literature on this channel as well, much of it devoted to trying to determine the reason for its non-selectivity. The TVGYG sequence has one mutation, to TVGDG, with the consequence that the upper two sites are replaced by a second aqueous vestibule. The open and closed structures have been determined at high resolution (36, 49, 50). In this channel, both Na^+^ and K^+^ maintain a symmetric configuration. The reason for the loss of selectivity has been variously interpreted in terms of small structural adjustments by a number of authors, without a general agreement (51-53). These citations are not comprehensive, but are sufficient to see that there is still not a general agreement as to how the NaK channel helps elucidate selectivity in selective channels.

A controversy that remains concerns the order of ions and water in SF. There are proponents of what Strong et al (24) refer to as “hard” and “soft” knock on mechanisms, where “hard” means that ions can be adjacent to each other in the SF (earlier referred to as “direct”), while “soft” would have a water molecule between ions.

Despite the extensive effort by multiple groups to understand ion selectivity, and the passage of ions through the SF, the subject still requires further work. Our calculation shows how an ion moves from the cavity in the K_v_1.2 channel into the lowest position in the SF, and compares, at an atomic level, the charges on the K^+^ ion and the neighboring solvating oxygens, to those when Na^+^ approaches the filter, and to those in a mutant in which threonine is replaced by serine. K_v_1.2 being larger, hence more difficult to understand, it has been less studied than KcsA. The calculations in the literature for the most part do not examine the transition from the cavity to the filter in atomic detail, or, as with MD simulations, there are serious problems that remain. They do note that the filter may be flexible, and some show a relation to inactivation, to ordered water, and sometimes to entropy. These calculations do not consider the reason serine is so rare, with threonine almost always the lowest residue in the SF. Some models are severely truncated, but done at QM level, and others are complete but done by MD with classical potentials. The KcsA channel pore differs from the voltage gated K_v_ channels, such as Shaker or K_V_1.2, not only in lacking a voltage sensor, but also in that the KcsA cavity contains less water, with the volume of the cavity being appreciably smaller. However, the SF is the same, and there is sufficient water even in KcsA to participate in solvating the ion at the entrance to the SF. De and coworkers (30) have used ONIOM 2-layer calculations, with the center of the filter in the QM section, to sort out the contributions of the individual amino acids in the filter sequence. They do consider hydration at the filter entrance, finding it endothermic, hence the entrance to be kinetics dependent. They do not consider the serine mutation. Channels with T→S mutations show somewhat reduced polarizability, and possibly reduced conductivity (52); Rossi et al attributed stronger binding of the K^+^ ion to the polarizability of the four extra methyl groups in the WT compared with the serine pore (54).

### Reasons for using quantum calculations

This said, there are a number of advantages to a QM calculation. The charges on the atoms, and the bond orders, can be determined. The energies, even though at 0 K, are more accurate, as exchange and correlation energy terms that have no classical analogue are included. These are larger than the room temperature value of k_B_T, and probably larger than the entropy terms that are lost. Also, the channel must go through tens of thousands of cycles in its lifetime. Most calculations do not test whether this is possible under the assumptions of that calculation. MD calculations, to the best of our knowledge, have not been shown to return to the starting conformation (we omit steered MD calculations here, which do not bear upon this point). We have been able to test one case for the stability of the calculation, using our QM calculations on a section of the voltage sensing domain (53), and found that it is possible to get back to the same minimum from different starting positions. Of course, one instance is not conclusive, but it does suggest that the QM results are not necessarily caught in local minima. The calculations presented here also show that the differences in energy that are found are at least reasonable; if there were serious problems with traps in local minima, we would expect large differences in the energy between different conformations, and the results should be inconsistent with the overall expectations. This is not found, although each minimization (optimization) is an entirely independent calculation. We believe this is significant evidence that the calculations avoid falling into inconsistent local minima. In our calculation, we use a large enough section of the protein that the interaction of the ion with the protein includes at least a second ring of atoms around the ion. Thus most of the surroundings contribution to the energy is included; Dixit and Asthagiri (43) have shown that the surroundings contribution is not large in general, and the fact that we have included at least all of the first shell of protein atoms and much of the second means that this calculation should be accurate, at least from this point of view. The more distant protein would be essentially the same for all ions, and all cases, removing one source of error when taking differences; errors of this type would cancel to a rather high degree of accuracy. All energy values used here are differences.

It is clear that there has been a huge amount of interest in the question of how ions traverse the pore of potassium channels, (as well as other channels, e.g., Na^+^, Ca^2+^, not discussed here). Each possible effect on the ion’s path has been considered, not always together. Most of the approaches either use classical calculations or simplified models. Taken together past work has helped in understanding aspects of the conductivity of the pore of potassium channels, but still leaves much more to be done. We present QM calculations here that show in detail how an ion enters the SF at the lower end of the SF. This too is a subset of the entire problem, but by including all relevant atoms in the region, examining a K_v_ channel which can be compared to the smaller KcsA channel which we had studied earlier, and paying full attention to the transfer of the ion from hydration in the cavity to cosolvation by the threonine at the beginning of the TVGYG sequence in the SF, all using QM calculations to insure the inclusion of the electrostatic, polarizability, and van der Waals effects, that have previously often been considered separately (55) as well as the exchange and correlation energy totally omitted in classical calculations. In some cases, we will see that electrostatics dominates, but it is worth making sure that all are included, as a difference of even 5 kJ (≈2 k_B_T, where T is assumed to be about room temperature) can be significant, and this much or more can result from non-electrostatic interactions. The water takes an interesting configuration with the ion in the cavity, which appears to be that which is seen in the X-ray structure. Oakes et al (56) in a new MD simulation of KcsA, including two mutations (T75A, T75C, where T75 corresponds to T374 in the K_v_1.2 channel) consider variations on a knock-on mechanism, with some conformations KWK (ion-water-ion) and some in which the potassium ions are directly apposed (KK). They do not, however, consider specific bonding or bond angles for hydrogen bonds. Their conclusions are similar to those of previous work, in which the KWK conformation is more common than KK; they have cavity water spacing in general agreement with X-ray electron density. They do not observe the “basket” of water first reported by Kariev et al for the KcsA pore (57), and found again in this work with the K_v_1.2 channel.

There has been considerable electrophysiological data in the older literature that is also very informative, especially now that calculations give atomic level results that should match the data, at least qualitatively, and actually almost quantitatively. Nimigean and Miller (58) studied the transmission of K^+^ through the channel, and concluded that there were barriers both in the channel cavity and in the selectivity filter. Our calculations show how this must be, in atomic detail. They also studied the transmission of other ions. Much earlier Hille (59) found that the ratio of permeability for K^+^ to NH_4_^+^ was 1.00:0.13; quantitative differences among Na^+^, K^+^ and NH_4_^+^ in the channel pore that appear in our calculations provide mechanistic insight. Our calculations on the T→S mutation (T374S in the 3Lut numbering) concern particularly the bottom of the SF; differences include hydrogen bonding and ion coordination, both in KcsA and K_v_1.2, with its additional water. It is known that this mutation remains selective for K^+^ (16), so that the most serious question is how the transfer from the cavity to the SF is arranged. The actual current for the mutation has not been reported, and we expect the current in the mutant to be reduced. Our calculated energy suggests that the transfer to the filter becomes more difficult by perhaps an order of magnitude in the mutant. We will show how the K^+^ ion is symmetric in the WT channel, while the channel with serines substituted for the four tyrosines becomes slightly asymmetric, with the ion moving towards a crevice that forms when shorter side chains of the serines leave four potential wells, instead of the single well for the WT. Our results agree, for the WT, with the structure, including in particular the water “basket” mentioned earlier; a very similar structure has been reported (as an antiprism) by Zhou and Mackinnon (3,4). We have also calculated the atomic charges which help in understanding the behavior of the ion, as well as the reason for the loss of symmetry. Our results in this sense include results beyond those of Rossi and coworkers (54), although we do not obtain thermodynamic quantities, as theirs do.

Many papers cite a “knock-on” mechanism, in which ions push the ion ahead in the transmission sequence forward. However, it is rarely clear why there should not be a “knock-back” mechanism, in which the ion ahead, which exerts the same force on the ion following as that ion exerts on the forward ion, pushes the following ion back. The channel is known to be a rectifier; why does it rectify? We show how the solvating oxygen groups have charges that pull the ions from the cavity into the filter. The ion is then held by the charges on the solvating oxygens, which may include the water ahead of the ion in the pore, so that the ion is not pushed back, and any repulsion with the forward ion serves to push that ion forward. Without determining the charges, the initial step into the SF is hard to understand. This still does not completely explain rectification. Most important, we find that a set of four waters behaves as the pawl in a ratchet and pawl mechanism that does not allow the ion to come back, accounting for the rectification implied by the knock-on mechanism. The results on Na^+^ ion show that there is no reason, in principle, for an ion to always move forward; Na^+^ actually takes a backward step.

## Methods

The system is truncated from a calculation with 870 atoms and no ions, as shown in Fig. 1A. Fig. 1B is an example of a starting configuration derived from that in Fig 1A, with 457 atoms, and shows the arrangement of the oxygens and two ions at the top of the cavity and the bottom of the selectivity filter, together with the surrounding protein atoms. In figure 1A five levels of oxygens are shown as large spheres. There are two K^+^ ions (gold) in Fig. 1B. The lowest level of oxygens (light blue) are in the cavity. In the case shown here the lowest K^+^ is in the cavity, just above this level of water molecules. Four water molecules (dark purple) above this level are hydrogen bonded to a level above in which the oxygens (small light purple spheres) are from the lowest threonines in the TVGYG sequence of the selectivity filter. These are the waters mentioned above, with their hydrogen bonds to the threonine oxygens, which form a “basket” similar to the structure found by Zhou and Mackinnon (4), which they described as an antiprism; the symmetry is less exact here, so we do not label it as an antiprism. There is a water molecule above the threonine oxygens (light blue sphere with two dark hydrogens). The second K^+^ above this is solvated by carbonyl protein backbone oxygens (light purple). This constitutes the S4 location in the selectivity filter The system in each case was optimized using HF/6-31G*, using Gaussian09 (60). The energy reported is calculated by single point calculations using B3LYP/6-31G**, which was also used for the NBO calculations (61, 62) that give the atomic charges. The calculation for Fig. 1A began from the X-ray structure (3Lut (63)), including 720 protein atoms from the pore, starting with the gate and extending to the second SF position (S3). 50 water molecules were added to this, for a total of 870 atoms, and the result was optimized. From this structure, 38 water molecules were deleted, leaving 12. 419 of the protein atoms were kept, the rest deleted; two ions were added, for a total of 457 atoms. This was the fundamental starting structure for the wild type (WT) cases (Fig 1B). With the T374S mutation, there were 12 fewer atoms, and with NH_4_^+^ as the ion, 8 more atoms. The calculations consisted of optimization of the upper section of the cavity, with the first two positions of the SF, generally designated S4 and S3, the optimization starting from the 3Lut structure. In each case, 52 atoms were frozen to prevent overall motion of the system; all the frozen atoms were protein backbone atoms that did not border the pore. The locations of the end of the serine chain in the initial structure were adjusted by hand.

**Fig. 1:**
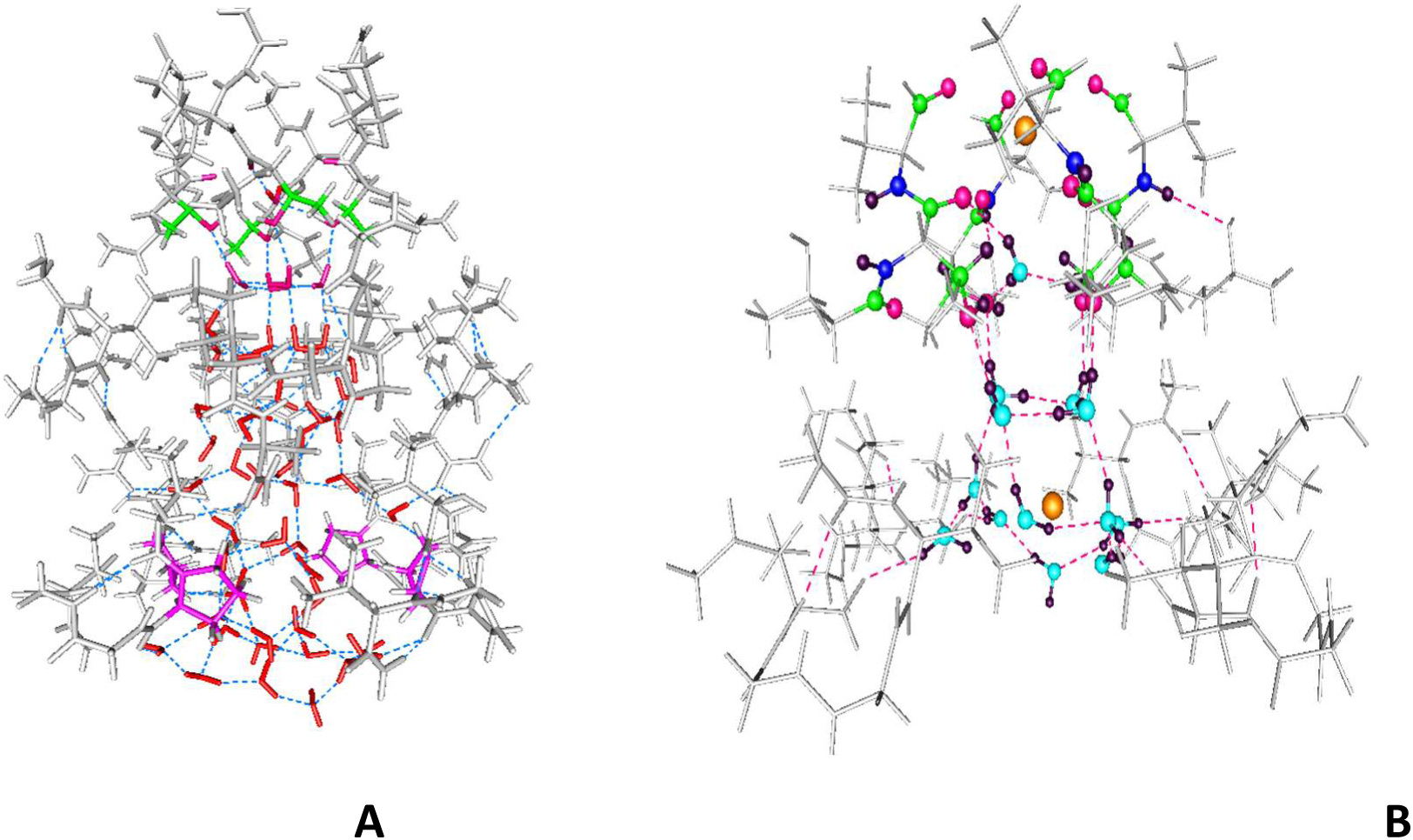
Fig. 1A shows the system with 870 atoms, but no K^+^, from which the starting conformation 1 (Figure 1B, for the K^+^ ion case) was derived.

### Definitions

*We use “conformation” to include the complete system, with both ions; “position” refers to the location of individual ions. We use “location” (or occasionally “position”, where the latter is unambiguous), to refer to parts of the protein, such as the threonine –OH, or the S3 or S4 locations in the SF. There are 4 “cases”: K*^*+*^*(WT); K*^*+*^ *(T374S); Na*^*+*^*(WT); NH*_*4*_^*+*^ *(WT). Each case was calculated in three conformations*.

The three conformations which were optimized were defined as follows:

*Conformation 1*, all cases: Initial configuration: one ion placed close to the S3 position from the X-ray structure, a water molecule in S4, and the other ion placed below the basket, in the cavity. This was then optimized. Figure 2 shows the position of the most relevant atoms for K^+^, Figure 3 for Na^+^, making it clear how different the K^+^ and Na^-+^ structures are after optimization. In each case, in conformation 1, the ion was started from a position on the pore axis symmetric with respect to the solvating oxygens. The lower ion was initially solvated by the basket water molecules for all cases. S3 is near the top of system as calculated, the pore being truncated one layer of atoms above this.

**Fig. 2:**
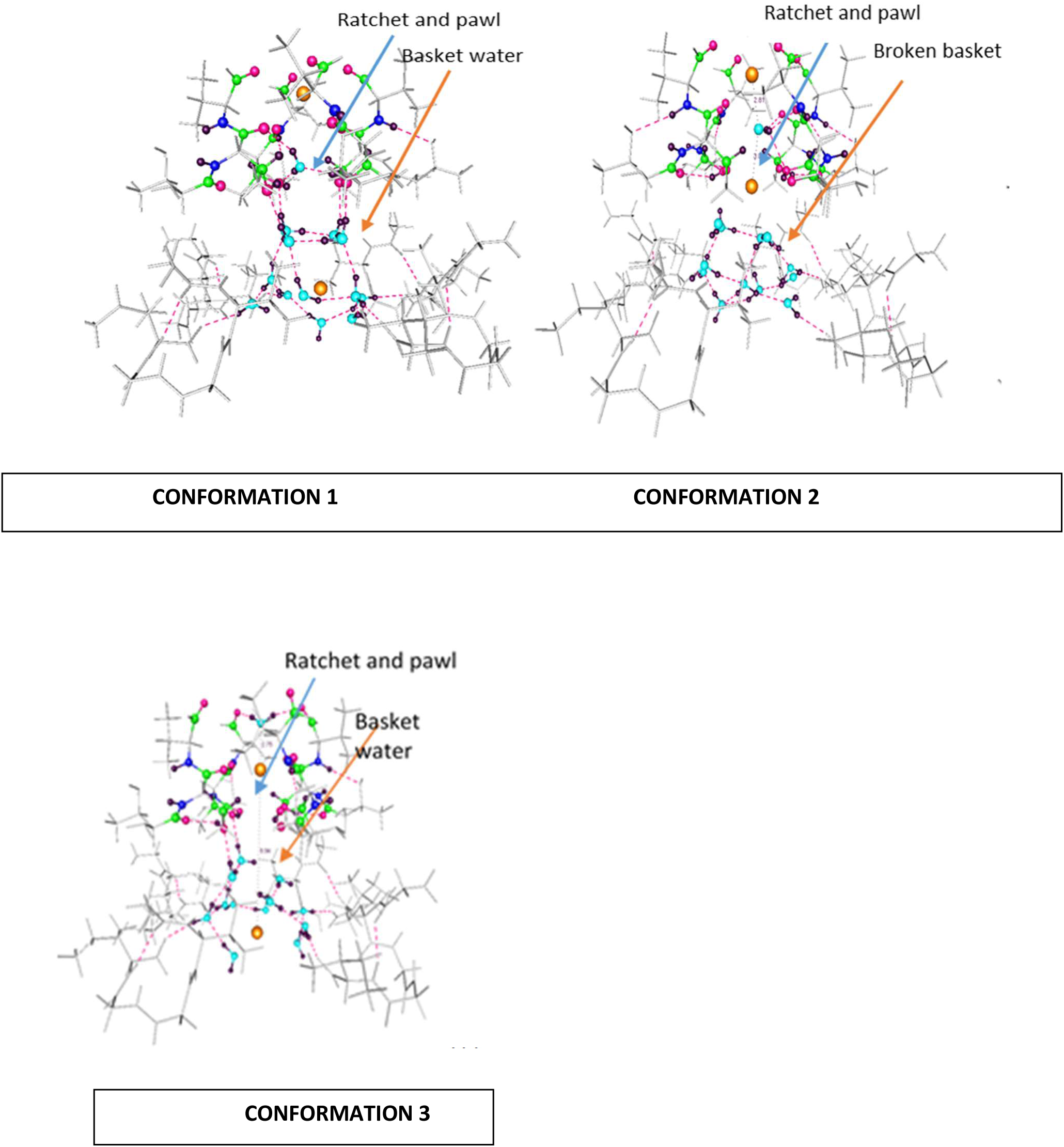
The three conformations with K^+^ in WT pore. Ions are large gold spheres. The basket of water is indicated for conformation 1 and for the partial recovery in conformation 3, while the broken basket that exists as the ion passes through is shown in conformation 2 (indicated by orange arrows). The level of the ratchet and pawl mechanism is indicated by the blue arrows; the relevant hydrogen atoms from the hydroxyls are shown in black (small spheres). C atoms, green; N, dark blue; solvating water O, light blue; carbonyl O atoms solvating the ion at bottom and top of S3, magenta; all other atoms, gray, including the 52 frozen atoms. The surrounding atoms of those relevant to the ion progress (gray) remain nearly constant throughout the calculation.

**Fig. 3:**
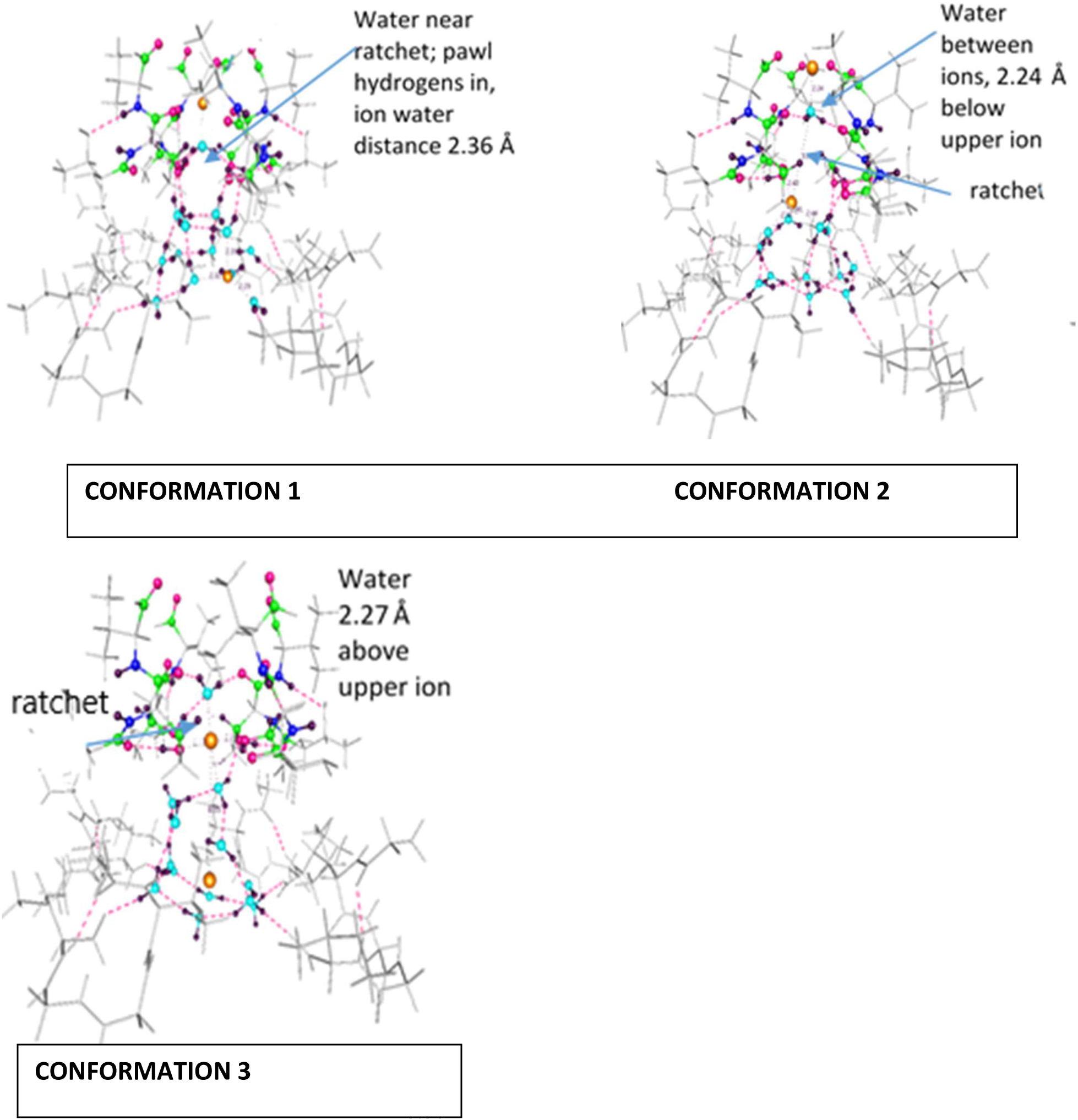
**Na^+^:** The corresponding Na^+^ conformations: Same colors and conditions as Fig. 2. Note the off axis position of the lower Na^+^ in conformations 1 and 2. In conformation 3, the Na^+^ has retreated below the S3 position, has a water above it, and waters below at hydrating distances, so that it cannot move. In conformation 1, the water designated as “near ratchet” is oriented with protons down, in contrast to all other cases, and the upper ion-water distance is 2.36 Å, much less than in the K^+^ case (3.97 Å), so it solvates Na^+^ strongly. The basket is intact. In conformation 2, with the lower ion still off-axis, and tied via electrostatic interactions to the pore wall, the upper ion is strongly solvated, and a water is only 2.24 Å below, thus tightly hydrating it; in going to conformation 3, the upper ion actually retreats, and a water *above* is 2.27 Å away, and tightly hydrogen bonded to the protein as well.

**Fig. 4:**
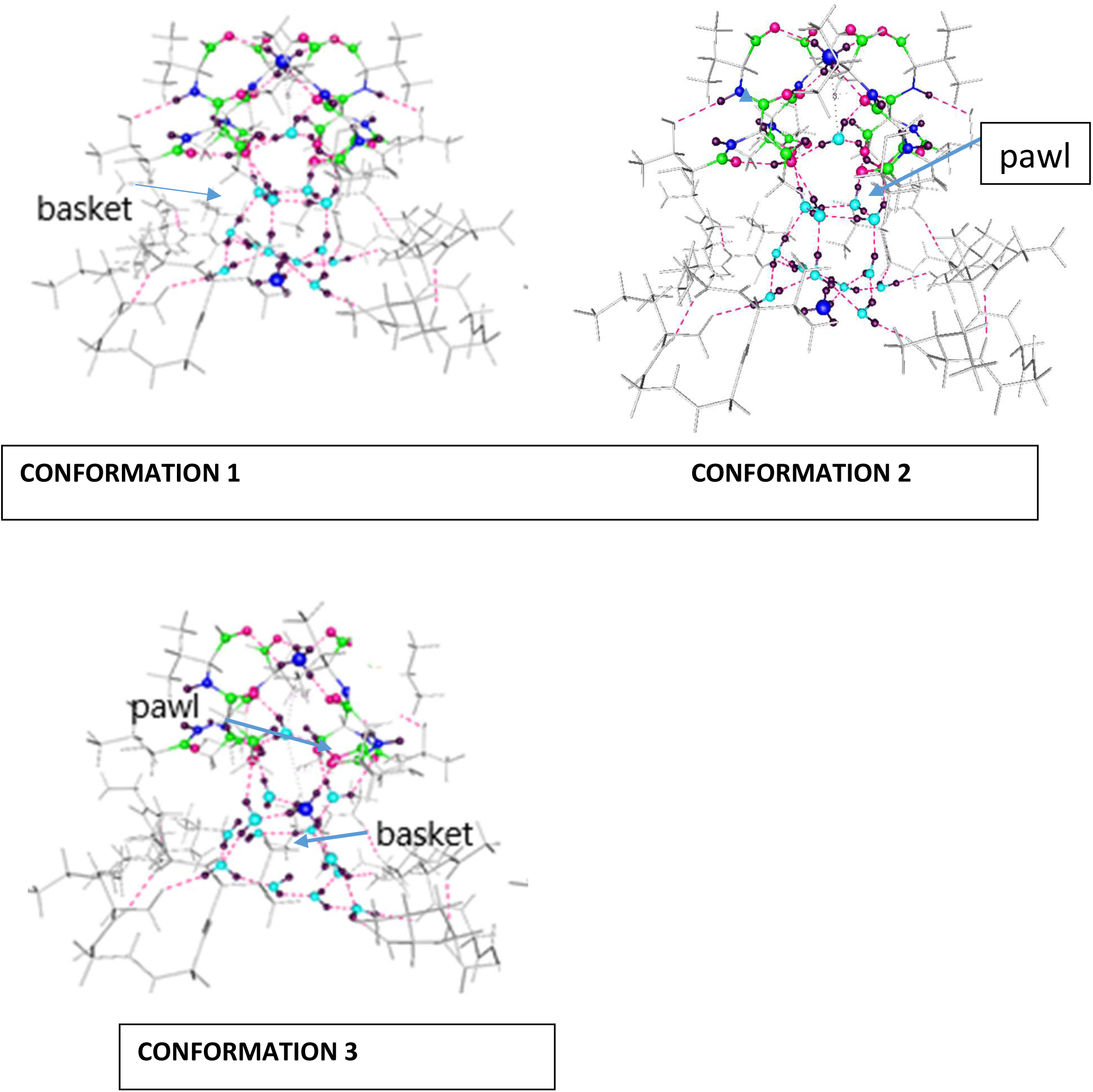
The NH_4_^+^ in the three conformations. The basket (conformations 2,3) and pawl (conformations 1,3) are indicated on the figures. The basket begins to reform in conformation 3. The NH_4_^+^ remains essentially on the axis, avoiding the asymmetry of the Na^+^.

**Fig. 5:**
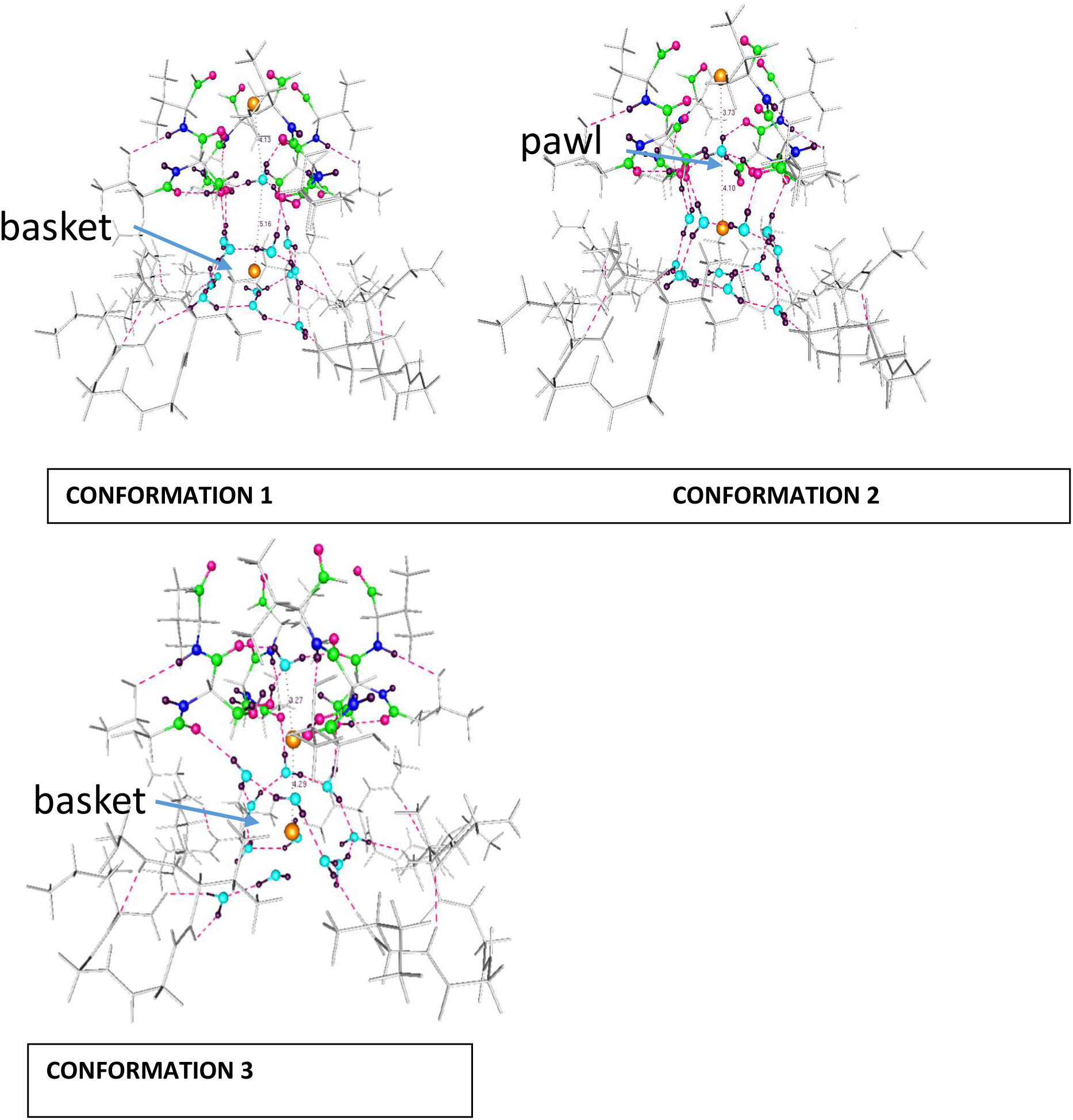
The three conformations for the T374S mutant. All color conventions as in Fig 2 and 3. The ions stay on axis; unlike Na^+^, symmetry is almost maintained. However, progress is impeded by the ratchet and pawl mechanism; some hydroxyls are folded into the ion path; see Table 5, and discussion.

C*onformation 2*, all cases: The calculation began from the optimized conformation found from conformation 1, except that the lower ion was moved up approximately 3 Å to a location between the basket and S4; it was optimized from this start.

*Conformation 3* started from the optimized conformation 2, with the ions moved such that one ion was below the basket, which is in all cases partially broken (see Figures) and the position of the other ion depended on the case, as the positions of the ions were not always the same after conformation 2 optimization. The point is to start with what would be the consequence of the upper ion moving up and out of the calculated range, the pore water below moving into this position, and a new ion coming up from below in the cavity. It would be a third step after the ions had gone through the first two conformations. The essential characteristic of this conformation for K^+^(WT) has one ion fairly near the S4 position, with water in the upper (S3) position. The NH_4_^+^ and mutant cases had also diverged enough after conformation 2 optimization to require specific description, which follows in the results section, and is illustrated in the figures. Except for K^+^ (WT) some asymmetry developed during optimization, which appears to have been especially important for Na^+^. NH_4_^+^ was asymmetric, but moved by a rather different mechanism, more like a water molecule in that it formed hydrogen bonds.

## RESULTS

*Conformation 1*: With the K^+^ ion, there was hardly any change in WT from the essentially X-ray position start. With the Na^+^ ion, however, the ion in S3 (upper ion) actually moved **down**, back towards the cavity, to a position in which it was held by four solvating groups, including the one pore water. Because the S2 location is not present, the interpretation is limited. The Na^+^ result is clearly different than the results with K^+^. For the K^+^ ion in the T374S mutant, the result in conformation 1 is the same as WT, as neither ion is near the serines. The NH_4_^+^ behaves somewhat like a charged water molecule; it can form hydrogen bonds, and seems to take the place of water in conformations 2 and 3. The charge drives it forward, however. We have not tabulated the results from conformation 1, in which the lower ion is below the basket in all cases, and the lower ion does not move.

The results from the optimizations are expressed quantitatively in Tables. Tables 1 and 2 correspond to conformations 2 and 3. Conformation 1 is not included, in which one ion is placed below the basket, as the K^+^ ion moved very little. There are two important results that begin to be understood from this conformation:

**TABLE 1.** Distances (Å)^+^ of ***lower*** ion to nearest 8 oxygen atoms in c**onformation 2**;

| → ++ | OH 1 | OH 2 | OH 3 | OH 4 | W 1 | W 2 | W 3 | W 4 |
| --- | --- | --- | --- | --- | --- | --- | --- | --- |
| $K^+ WT$ | **2.68** | **2.79** | **2.82** | **2.87** | **2.88** | **2.93** | 4.03 | 4.07 |
| $Na^+ WT$ | **2.40** | **2.65** | 3.93 | 4.02 | **2.30** | **2.40** | **2.44** | 3.75 |
| $NH_4^+ WT^*$ | **2.96** | 3.80 | 4.64 | 5.15 | **2.85** | **2.86** | **2.93** | **3.07** |

<sup>+</sup> The distances apply to the final position of the lower ion, starting (pre-optimization) between the basket and S4; the upper ion starts in S3
++ T374 OH first four columns, basket water oxygens, last four columns
*\*NH<sub>4</sub><sup>+</sup> the distance is given to the N atom. Boldface distances show solvating oxygens*

**TABLE 2.** Distances (Å) of the ***upper*** ion, now in position S3, to the nearest 8 oxygen atoms, in **conformation 3**.

|  | +NCO<br>1 | NCO 2 | NCO 3 | NCO 4 | OH 1 | OH 2 | OH 3 | OH 4 |
| --- | --- | --- | --- | --- | --- | --- | --- | --- |
| *K<sup>+</sup> WT* | **2.89** | **3.08** | **2.86** | **2.85** | 3.95 | **3.29** | **3.22** | 4.70 |
| *Na<sup>+</sup> WT* | 4.81 | 4.53 | 4.49 | 3.73 | **2.88\*** | **2.29\*** | **2.90\*** | **2.47\*** |
| *NH<sub>4</sub><sup>+</sup> WT\*\** | 5.27 | 4.09 | 5.28 | 5.28 | **2.81** | **3.08** | **2.88** | **2.79** |
| *K<sup>+</sup> T374S* | 5.48 | 5.29 | 4.63 | 5.12 | **2.78** | **2.79** | **2.85** | **2.75** |

+ Distances to T374 carbonyl oxygens, except for NH<sub>4</sub><sup>+</sup>
\* Distances to basket waters below the SF; Na<sup>+</sup> has failed to advance
\*\*NH<sub>4</sub><sup>+</sup>: the distance is given to the N atom. One basket water remains in its solvation shell
For the mutant with K<sup>+</sup> two basket waters remain in its solvation shell, in addition to the T374 OHs

### Selectivity

The *upper* Na^+^ ion, starting in S3, moves **down**, as noted above. The K^+^ ion starting in this position holds its place; it cannot move up, as the section we calculate does not have S2, but it does not move down, as Na^+^ does; the Na^+^ is held by three waters in the SF, between the ions, at a distance of 2.36 Å. A fourth oxygen from threonine completes the solvation of this ion. This suggests at least part of the reason for the difficulty Na^+^ has in making progress toward the extracellular surface in going from conformation 1 to conformation 2 (but see also large positive ΔE, Table 3). Specifically, the energy that is required to break the solvation occasioned by the short 2.36 Å oxygen-ion distance is great enough for the ion to be effectively trapped. The short distance is possible because the ion moves off axis; a symmetrically placed ion, like K^+^, is sufficiently distant from four oxygens that it is not held as tightly. This appears to have a major role in selectivity, because the short ion – oxygen distances hold the ion tightly.

**TABLE 3.**
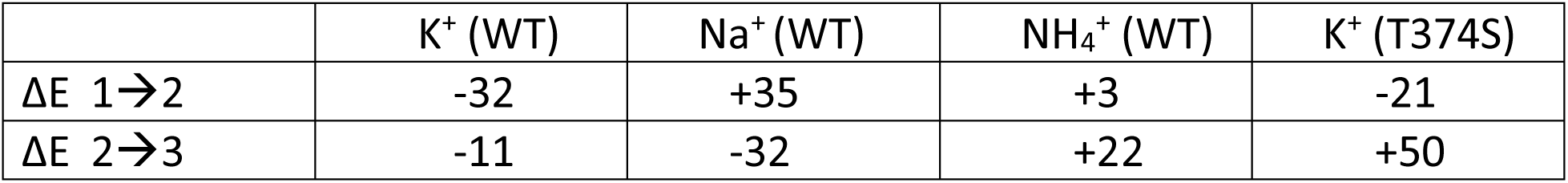
Total system energy change (kJ mol^-1^) for the transfer of the ion from conformation 1 to 2, and from 2 to 3

### Rectification

There is one other important difference in the arrangement of both the T373 backbone carbonyl oxygen and T374 –OH groups. These oxygens can either point into the pore or flip back away from the pore. With K^+^ the –OH forms a hydrogen bond with the water in the S4 position for three out of four –OHs. One flips to form an H-bond with a T373 carbonyl. With Na^+^ three OH flip, while one forms an H-bond with the S4 water. However, as the lower ion moves up, the K^+^ WT case shows these –OHs all flipping out of the way, and this is the only case in which this happens. See the discussion of the O – O – H angle below as a ratchet and pawl (not a bond angle: the first O is from a carbonyl). Rectification must also be explained, because the force exerted on an ion rising into the selectivity filter is the same as the force exerted by the ion to push the ion in the selectivity filter forward. Hence, an ion could be pushed back, reversing the current. This mechanism appears to account for the fact that ions must move in one direction.

The lower ion described in Table 1 (conformation 2) moves up, and becomes the upper ion in conformation 3; the lower ion in conformation 3 is below the basket. In effect, this assumes the former lower ion in going from conformation 2 to conformation 3 has been “knocked-on”, becoming the upper ion, while a new ion enters the system as the new lower ion. The upper ion in conformation 2 has moved into the upper part of the SF, therefore no longer within the computed section. (as quantum particles, the ions are not actually distinguishable, so describing the ions as though they could be labeled assumes a somewhat classical viewpoint, but it should be clear that the overall conformations and transitions between them are being described.) Table 1 shows that only K^+^ in the WT has symmetric distances to the next level up, defined by the four threonine –OH distances. The total number of solvating oxygens (boldface) for all cases is 5 or 6, except in the mutant in which only 4 oxygens are within the appropriate distance to solvate the ion. The K^+^ in the WT has its solvation symmetric in this plane. In the other three cases, only one or two of the –OH oxygens are within solvating distance, making the ion take an asymmetric position in this plane, or just below it. The K^+^(WT) case has the ion breaking the basket, so that the distances are unequal in that plane, but the ion remains essentially on the pore axis. This is not true of any of the other cases. In the mutant, the ion makes a little progress, while the Na^+^ case has the ion asymmetric, and close to its initial position vertically, unlike the K^+^ ion. In other words, only the K^+^ ion makes forward progress into the SF. Alternatively, one can think of a competition between the basket water and the threonine –OH, which is won by the threonine –OH only for the K^+^(WT) case. The NH_4_^+^ ion hardly moves from its initial position, remaining largely solvated by the basket waters. Its tetrahedron is oriented with three H atoms down, one up.

#### Boldface distances show solvating oxygens

Table 2 gives distances for the same ion for which the distances were given in Table 1, but with the ion in a different position. It gives the corresponding distances for the upper ion in the transition from the threonine oxygen plane to what is essentially the S3 position. The distances in the first four columns are to the oxygens of the backbone carbonyl groups.

The lower ion in this conformation starts in the cavity below the basket. Examination of the solvation shells shows Na^+^ stuck with strong solvation by basket water from the cavity, and failure to advance to the SF. The NH_4_^+^ and K^+^ mutant advance, but not completely. This set of distances shows a clear difference between Na^+^ and K^+^ in the pore. Only K^+^ (WT) has strong solvation from the carbonyl oxygens, and no solvation from basket water. None of the other cases has the ions optimizing close to the carbonyls.

Table 3 shows the transition energies between the various structures, as the differences in total energy of the system with the different ions, with the mutation, in kJ mol^-1^. For comparison, k_B_T is approximately 2.5 kJ mol^-1^, so 20 kJ mol^-1^ in the Table corresponds to a factor of exp(8) ≈ 3000; that is, a barrier of 20 kJ mol^-1^ would slow the ion by a factor of roughly 3000.

Table 3 shows the change in energy is negative both for a K^+^ ion to go from the cavity to the plane at the bottom of the lowest position in the SF (S4), and then on to the top of S4. All other cases show at least one large energy increase, which would suggest that there is a substantial barrier to the progress of the ion, but this must be understood with caution. Lack of an energy barrier is not sufficient to allow the ion to progress; for one thing, for the mutant, and especially for Na^+^, the ion goes backward instead of to the top of S4. The energy goes down, but this does not necessarily mean the ion goes forward. Energy can go down if the next conformation is of lower energy, with the ion going backward as with Na^+^ going from conformation 2 to 3. Also, there is a ratchet and pawl mechanism that can block progress, and appears to do so in every case except K^+^ (WT). We will discuss the ratchet and pawl mechanism for insuring K^+^ goes forward below. Unlike Na^+^, the K^+^ at the S3 position moves forward out of the computed region; it is replaced by a water, which moves up from below. This is the only case in which there is no barrier to either step, either energetically or through the ratchet and pawl mechanism (Table 5). Na^+^ and NH_4_^+^ behave quite differently from K^+^ and each other. NH_4_^+^ has a substantial energy barrier in the last step, but, as we see below, behaves more like a charged water molecule than like metal ions, and it can move up, albeit not as well as K^+^; its conductivity is about 0.13 that of K^+^ (59). The NH_4_^+^ conductivity mechanism is different from that of the metal ions, and it is possible to understand how this ion moves forward in spite of asymmetry, or the O – O – H angles, by examining the selective hydrogen bonding. Finally, the T374S mutant shows the ion essentially unable to move past the first step. In this case, the result is consistent with the one large energy barrier. The ion in this case breaks the water basket below the SF, and retreats, somewhat as Na^+^ does in the conformation 2 to conformation 3 transition. The hydration of the ion in the first step switches from the basket to the threonine hydroxyls (except that in the T374S case, it is serine hydroxyls). We know from experimental results that the selectivity for K^+^ over Na^+^ is large, that it is less for NH_4_^+^, and that the serine mutant allows K^+^ to pass, but not as well as the WT.

**TABLE 4.** Ion charge for the relevant cases*.

|  | K<sup>+</sup> (WT) | Na<sup>+</sup> (WT) | NH<sub>4</sub><sup>+</sup> (WT) | K<sup>+</sup> (T374S) |
| --- | --- | --- | --- | --- |
| Conformation 2 | **0.80 / 0.76 | 0.79 / 0.75 | 0.92 / 0.84 | 0.78 / 0.81 |
| Conformation 3 | 0.75 / 0.82 | 0.74 / 0.73 | 0.84 / 0.83 | 0.77 / 0.78 |

\* Recall that conformation 3 has the ion which was originally the lower ion rising to the S4 position of the SF; conformation 2 has the lower ion in the position in which it optimizes in the first transition from below the basket, as shown in the Figures. However, for Na<sup>+</sup> the ion moves backward, so the ion position in conformation 3 differs from that of K<sup>+</sup> (see Figures). However, this leads to only limited difference in charge transfer
\*\* The left hand number in each cell is the charge on the ion in the upper position (e.g., for K<sup>+</sup>(WT), in S3 for conformation 2); the right hand number is the charge on the ion in the lower position, in the conformations shown. The NH<sub>4</sub><sup>+</sup> charge is the sum of charges on all five atoms.

**TABLE 5.** Number of angles (O(T373 –C=O) – O(T374 –OH) – H(T374 –OH) in 3 ranges

| | $K^+$ (WT) | $Na^+$ | $NH_4^+$ | $K^+$ (T374S) |
| --- | --- | --- | --- | --- |
| Angle* | <25°/>90°/M+ | <25°/>90°/M | <25°/>90°/M | <25°/>90°/M |
| Conformation 1 | 1/3/0 | 2/2/0 | 1/3/0 | 2/2/0 |
| Conformation 2 | 4/0/0 | 2/1/1(35°) | 1/3/0 | 1/2/1(53°) |
| Conformation 3 | 3/0/1(35°) | 3/1/0 | 3/1/0 | 2/2/0 |

\* O(T373 –C=O) – O(T374 –OH) – H(T374 –OH)
+ angle: $25^\circ < M < 90^\circ$

### Error limits

These energy results are qualitatively consistent with the observed behavior, even if the energy differences, especially for the mutant second step, seem somewhat too large. Because the absolute values of all calculations are those corresponding to the same outer section of protein atoms in the calculation, with presumably the same errors due to truncation of the system, the differences are likely to be more accurate than the absolute values, due to cancellation of errors in nearly identical systems. Interaction of the center of the system with the boundary may differ in the several cases by enough to account for only a small error. An error of >20 kJ mol^-1^ would be required to invalidate any conclusion stated here (the mutant conformation 2→3 transition may be so large as to be an exception). The most important difference is that between K^+^ and Na^+^. From conformation 1 to conformation 2, it is 67 kJ mol^-1^, which seems reasonable. While the errors are almost certainly not large enough to affect conclusions, they might be on the order of 10 kJ, and qualitatively the same results would be obtained for errors up to 20 kJ. This said, selectivity can be better understood by examining structure than from these energy values; they are, however, qualitatively consistent

There is also the fact that these are 0 K energies, and they may not all have the same heat capacities, hence the ΔG values at room temperature may differ somewhat from the 0 K ΔE. However, the differences should be small, as the structures are similar except for the ion’s positions and immediate solvation shells. Unfortunately, it is not possible to use our present computer resources to obtain these values for two reasons: thermodynamic quantities are computed assuming a normal mode analysis, which, with a reasonable basis set requires too much memory, and more fundamentally, a normal mode analysis is likely to be in error. The non-linearities occasioned by the coupling of modes is already enough to produce errors larger than the differences between the different cases. That the order of magnitude is reasonable is important to assure us that the calculations are not badly in error; the differences are between completely independent calculations. If the calculations were inconsistent, they would show this in huge values of energy differences, but that does not happen (Table 3); if inappropriate energy minima were reached by different calculations, this kind of error would be expected; each calculation requires a separate minimization, so if significantly different minima were found by the optimization, that would be apparent. We are therefore justified in taking structural results as basically reliable.

It is difficult to separate the effects of electrostatics, van der Waals terms, and of charge transfer back to the ions; this may not even be a meaningful statement, as the charge transfer leads to electrostatic consequences. The strictly quantum terms (exchange and correlation energy) can be determined separately, but not easily interpreted; they are smaller than the electrostatic terms, but not negligible compared to k_B_T, which is relevant. Almost certainly, the van der Waals terms are smaller than the electrostatic terms, but they may not be negligible, as the differences may be of the order of a few kJ mol^-1^; again comparable or slightly larger than k_B_T. We observe (Table 4) that the K^+^ and Na^+^ ions both have charge of about 0.78 ± 0.04 *q* (*q* is the electronic charge) in most conformations and positions. This seems to remain roughly constant in the conformations that we calculate here. The total charge on the NH_4_^+^ ion (lower position) is somewhat larger. The charge for the K^+^ ion (lower position) in the mutant is still approximately 0.81 in conformation 2. We can see that there is some charge transfer, with a fraction of an electron moving to the ion, but that there is not a great deal of difference as the ion moves from one position to another. Electrostatics remains the largest term in driving ions forward, however, especially for K^+^ in the WT channel because of the charges on the protein groups. At the top of the S4 SF position, the four atoms at the corners of the positions (H-N-C=O of T374) have charges of −0.161 ±0.05; the bottom corners of the S4 position (H-C-O-H, also of T374), +0.058 ±0.08. Summing the four corners, there is then a net charge difference of 0.876 (average) driving the ion upward. The thickness of this site is approximately 3 Å, the diagonal 6 Å(top), 5.3 Å bottom). The consequent field is sufficient to drive the ion from the bottom to the top of this section, helping in the rectification. However, there is another component in the rectification, the ratchet and pawl mechanism that we have referred to earlier.

### Ratchet and Pawl

*Hydroxyl Rotation Is a Key To Rectification:* There is one arrangement of three atoms that is critical for the progress of the ion through the selectivity filter: O(T373 –C=O) – O(T374 –OH) – H(T374 –OH). Using the T373 carbonyl oxygen as reference, the T374 side chain OH can flip its hydrogen into the ion path, or back away from it.

Table 5 shows that the T374 hydroxyl proton either bends back out of the pore pathway (the O(T373 –C=O) – O(T374 –OH) – H(T374 –OH) angle is usually ≈18°, always < 25°), or rotates into the path (angle >90°); of the 48 angles in the Table, only 3 take intermediate values, two of 35°, one of 53°, as shown. While the great majority of angles shown as <25° are actually ≈18°, there are two in conformation 3, K^+^ (WT), that are larger, ≈ 24°. The essential point is that angles <25° do not obstruct the path of the ion, while angles >90° do. Only K^+^(WT) has no obstruction going from conformation 1 to conformations 2 and 3. On the other hand, the conformation 1 angles show that the ion cannot move backward. This set of angles in the three conformations makes this section of the channel a rectifier. As the ion moves to conformation 2, the hydroxyls flip back out of the way, allowing the next ion to move forward. In effect, the hydroxyls act as a pawl in a linear ratchet and pawl mechanism. Combining Table 5 and the electrostatic charges of the S4 cube allows us to see how an ion moves forward. There is a charge difference going from the bottom plane of the S4 position to the top plane, in total, as noted above, averaging 0.876 q, spread out over approximately 60 Å^2^ and separated by roughly 4 Å. This constitutes the force driving the ion forward, analogous to the crank in a mechanical ratchet and pawl. Here, the molecular equivalent reorients with each passing K^+^, allowing it to pass. The knock-on mechanism relies on electrostatic interaction between ions, but the force pushing back the rising ion equals the force pushing the upper ion forward, so that we must account for the reason that the pair of ions always moves forward through the selectivity filter. Our calculations are not sufficient to give the details of the return to the initial configuration from the final step in which the upper ion moves ahead, but it is not hard to see qualitatively how the next ion, moving in below the basket, could restore the initial configuration.

For all the cases other than K^+^ (WT), the hydroxyls are not entirely cleared for the transition from conformation 1 to 2. The obstructions are not absolute, and do not alone account for selectivity, but do suggest that the channel is most easily traversed by K^+^ in the WT channel. The ratchet and pawl mechanism in the other cases interferes with the progress of the ion by having the pawl partially intrude into the ion path ahead of the ion, rather than behind.

We next need to compare the ion-pore interactions with specific detail, including the protein.

#### 1) K^+^ and Na^+^

We have seen in Table 3 that the K^+^ ion goes downhill in energy in both transferring from conformation 1 to 2 and from 2 to 3. The angle formed by the upper ion/water oxygen/lower ion is almost the same for both ions (167° for Na^+^, 162° for K^+^), so in both cases the pore is slightly bent. However, there is a substantial barrier for Na^+^ in going from conformation 1 to 2, and as we noted it retreats instead of moving forward. The water molecule in S4 rotates as the ion passes.

With K^+^ in conformation 2, Table 5 shows when the hydrogen of the T374 hydroxyl is bent back out of the way of the ion, or remains in the ion path.

A. K^+^ (WT): *Conformation 1*. The hydrogen forming the hydrogen bond (T373 carbonyl—T374 –OH) forms three such bonds in conformation 1 in the WT, with K^+^. In conformation 1 the ion is out of the way of the hydroxyl, so this is not a problem for conduction. Only one out of four hydrogens is bent back. *Conformation 2*: All four H from the T374 –OH are bent back (average O – O - H angle ≈18°). Now there are no hydrogen bonds between the threonine and the pore SF water, leaving a clear path for K^+^. The plane defined by the central water molecule is almost orthogonal to the pore axis (water plane angle to pore axis = 98°), (*cf*. Na^+^ for which the corresponding angle is 116°). With the water tipped down in this manner, the hydrogens are positioned to make hydrogen bonds with the T374 carbonyl oxygens, while with the K^+^ the hydrogens are not able to make such hydrogen bonds. *Conformation 3*: Essentially as in conformation 2, no interfering hydrogen bonds; the O – O - H angles are a little larger, about 24°. The distance from the upper ion to the water oxygen is 2.81 Å, meaning that they are closely associated; the distance to the lower ion, 3.08 Å, is enough larger that the association is appreciably weaker. In conformation 3, the water-ion distance is down to 2.75 Å, but this is water to the next ion, which has jumped up through the entire S4 position, with the four O – O – H angles we have been discussing all bent back out of the way. The ion that was above the water in conformation 2 has moved to the upper part of the SF, out of the calculated section.
B. Na^+^ (Fig. 3): *Conformation 1*: There are two obvious differences with K^+^ in conformation 1. First the lower Na^+^ ion is off axis. Second, there is a water molecule in the pore between the ions. In the Na^+^ case, it hydrates the upper ion rather strongly (distance to oxygen, 2.36 Å), and the hydrogens are below the oxygen. This is the only case in which the water takes this orientation, blocking the pore and interacting with the ratchet hydroxyls (three of the T374 residues). With K^+^ the distance to the upper ion is 3.97 Å, too great to contribute to the hydration of the ion. The lower Na^+^ ion interacts primarily with cavity water, below the basket. *Conformation 2*: Asymmetry, and one interfering H bond, remain and make it difficult to advance, as the ion’s positive charge is close to the negatively charged oxygen on one side of the pore (Table 1). The combination of the asymmetry in the Na^+^ - O distances, with the O – O – H angles shown in Table 5 adding somewhat to the effect, helps explain the barrier as shown in the energy increase in going from conformation 1 to conformation 2 (Table 3). Thompson et al found an X-ray conformation for KcsA with Na^+^ very much like what we calculated, save that they did not find the asymmetry. It is not clear that they allowed for the possibility of asymmetry; the placement of the ions and water is otherwise very similar to our calculation for conformation 2; they also found a large barrier for the ion to enter the SF, in agreement with our result for going from conformation 1 to conformation 2 (Table 3) (64). *Conformation 3*: This is the only case in which the ion went *backward* after conformation 2, which accounts for the negative energy for the transition (Table 3); it is not the case that the negative energy indicates lack of a barrier to forward motion, but lack of a barrier to backward motion. The ion is blocked in going from 1 to 2, so overall there is no progress for the ion in the SF. *Summary:* The location of the hydrogen bonds partially determines whether the ion can progress through the SF but it is mainly the asymmetry and, secondarily, the O – O – H bonds that constitute the pawl for the ratchet that impede forward motion (Table 5), so the ion is blocked by the hydrogens that remain in the path. For K^+^, but not for Na^+^, the pawl - O – O – H bonds bend all the hydrogens out of the path of the ion in going from conformation 1 to 2, and from conformation 2 to 3.

There is another major difference between the ions. We have already discussed conformations 1 and 2. Consider the water molecule that separates the ions along the pore (the water in the ion – water – ion sequence). We just saw that in conformation 2, with K^+^, the upper ion – pore water oxygen distance is 2.81Å, lower ion to the same oxygen, 3.08 Å. For Na^+^ the corresponding distances are 2.24 Å for the upper ion, and a fairly huge 5.04 Å for the lower ion. It is possible that the energy required to separate this water from the ion is responsible for much of the large barrier shown in Table 3 for Na^+^ in passing from conformation 1 to 2. Once one gets to conformation 2, the upper Na^+^ ion is tightly hydrated, while the other ion is completely separated, and held by waters below, approximately in the basket location. These four conformation 2 waters have Na^+^ ion-water distances (d) 2.30 ≤ d ≤ 2.44 Å, so that they hydrate the ion well (recall the 2.36 Å distances in conformation 1), but, being also hydrogen bonded to neighboring protein groups, also anchor it in conformation 2. Since our conformation 3 assumed the K^+^ ion in the upper position, S3, had moved up, the inability of the lower Na^+^ ion to move might not show up clearly in the energy computation. K^+^ has transferred from hydration below to solvation by the T374 carbonyls above, and with distances to neighboring oxygens that suggest that the ion is able to continue to transfer almost exactly on the pore axis, while Na^+^ is trapped. In other words, moving to conformation 3 for Na^+^ is not progress.

This result can be looked at in a slightly different manner that reinforces the point. There are hydrogen bonds between the T373 carbonyl (this is the threonine below the beginning of the filter proper at T374) and the T374 hydroxyl. The asymmetry noted for Na^+^, but not K^+^, can also be understood from ion to hydrating water distances, especially in conformation 3.

Conformation 2: For K^+^ the hydrating water distances are 2.72±0.07 Å, for Na^+^ 2.80 ± 0.20 Å, with a range from 2.68 to 3.00 Å; the Na^+^ appears to twist the pore slightly (see Fig. 3).

Conformation 3: the distance for K^+^ is 2.78 ± 0.03 Å, while for Na^+^, the corresponding distances are 2.79 ± 0.29, ranging up to 3.08 Å. K^+^ hydration has the water molecules an order of magnitude more tightly grouped (i.e., symmetric) than does Na^+^. The overall structure of the pore is not greatly changed, but the symmetry is somewhat broken when Na^+^ is in the pore. Nimigean and Miller showed that it takes about 200 mV to drive the Na^+^ ion through the KcsA potassium selective SF (62), much more than physiological potentials. The Na^+^ ion is also solvated differently from K^+^ in bulk solution, which may be expressed through the stronger electric field for the smaller ion, in addition to steric consequences of the small size difference; the distances above suggest that both effects play a role. The differences between Na^+^ and K^+^ can be seen by comparing the conformation 2 figures.

#### 2) NH_4_^+^

*Conformation 1:* The most remarkable thing about conformation 1 is that it is unremarkable; the ions take positions similar to those taken by K^+^. Conformations 2 and 3 are different.

*Conformations 2 and 3:* The NH_4_^+^ ion differs from the other two ions in a rather obvious way, as it has five atoms, and stretches therefore over a distance which allows different hydrogen bonding partners. While this ion is less well able to traverse the pore than K^+^ (about 1/8 the rate of K^+^), it is much better able to do so than Na^+^. Also, there is a difference in charge transfer (net charge on the entire ion is about 0.84 in most cases, larger than for the metal ions) as well as the fact that it does form hydrogen bonds, rather than only electrostatic and charge transfer interactions as with the K^+^ and Na^+^ ions. The angles in Table 5 do not look promising, as three are large, and thus obstruct the path in conformation 2. However, NH_4_^+^ can rotate, so that its relation to the progress of the ion is not as direct as with the metal ions. While it appears that the decrease in the number of unfavorable hydrogen bonds (Table 5) in going from conformation 2 to conformation 3 should help, the barrier gets larger (Table 3). We can suggest that the rearrangement requires more energy, so that a larger barrier results. The figures help in understanding how this ion differs from Na^+^ and K^+^; the comparison turns out to be complex. In conformation 2, the lower NH_4_^+^ has inserted itself into what remains of the now asymmetric basket, replacing one water, and forming hydrogen bonds in its place. By replacing water, it appears that it can work its way through the selectivity filter, going off axis, but not in the way Na^+^ does, so it can produce a reasonable ionic current, albeit not as large as K^+^.

#### 3) K^+^ (T374S mutation)

*All conformations:* Here the difference with WT is a consequence of the difficulty the ion has in charge transfer and electrostatic interactions with the serine hydroxyls, which are a little more than 1 Å more distant in each direction than the threonine hydroxyls.

However, again the same O – O – H angles turn out to be the main issue. Two out the four hydrogens are bent back, but two are in the path of the ion. This is true in all three conformations. The key O – O - H angles do not allow free passage. The consequence is a major blockage in transferring from conformation 1 to 2. By the time the ion reaches conformation 3, it has broken through the barrier and reached a state somewhat similar to the ion in the WT channel.

Going back to conformation 1, we find that the H-bonds are only slightly asymmetric: two are similar to those in the WT, but one is with an –OH from T374, one with a –C=O from T373. As a consequence, the basket is distorted, impeding the ion’s progress. The distance from the lower ion to the S4 water is 5.16 Å, from the S4 water to the upper ion 4.13 Å, so that the solvation of the ion by these waters is weak to non-existent; the occupancy of the SF is distorted, and, with two O – O - H hydrogens blocking the ion, the path is difficult. As shown in Table 1, the lower ion is solvated by three basket waters (the basket is now also distorted).

### Summary

The results with the T374S mutant support the basic lesson of the WT, that the position of hydroxyl groups, especially those on T374, and their hydrogen bonding, determines whether the ion has a clear path to move forward in the SF.

### Electrostatics of the lower transition

We saw earlier that the sum of the charges on the atoms in the plane above the S4 plane is more negative than those in the plane below, the difference being 876 mV. Therefore, this requirement is fulfilled. We can only postulate at this point that the charges above this remain approximately the same all the way to S0, the final position in the SF, so that the ion can exit to the extracellular medium with no gradient needed; that computation has not yet been completed. However, we note that the carbonyls are very similar to those in the upper S4 plane, so it is reasonable to assign roughly the same charges to them. If so, this, combined with the driving force and the ratchet and pawl in the lower section of the SF, would explain how the knock-on mechanism works in a forward only direction. From this, one might guess a mechanism by which K_ir_ channels have opposite rectification, although it is not clear where in the SF this would be; it is not necessary that it be the first or last position in the SF. It does suggest that a different, or differently oriented, mechanism, must be involved. This remains for future work.

## DISCUSSION

The transition of the ion from the cavity to the selectivity filter is complex. It requires that the ion be properly solvated in both the cavity and the SF, but quite differently. The exchange of solvation, with preservation of symmetry, works for K^+^ but not for Na^+^, and rather differently for NH_4_^+^, which is itself not spherically symmetrical, so that its orientation matters. The cavity → SF ion transfer appears to be responsible for much of selectivity. Each ion is solvated differently, both in the cavity and in the selectivity filter, and the transfer between the two locations requires that the ion be able to remain on the pore axis in order to continue through the filter. We can draw four lessons from our results: 1) not all ions are solvated equally in the transfer into the SF. 2) Threonine is the right size to insure the proper solvation of K^+^ but not Na^+^. 3) by examining the charges on the solvating groups, we understand the importance of electrostatic forces in moving the ions into the first position of the SF. 4) the orientation of water with respect to the pore axis and the protein is important in producing rectification. This is not a direct knock-on transition; such a mechanism would in any case require a trap for the lower ion, so that it did not become a knock-back mechanism, in which the upper ion pushes the lower ion back down. The ratchet and pawl part of the mechanism requires water between the ions. 5) Finally, there is a role of symmetry in the transition into the SF from the cavity, with part of the selectivity against Na^+^ coming from the slightly asymmetric placement of the ion, which increases local electrostatic interaction. The shorter serine side chain in the mutant disrupts the proper hydration, and produces some asymmetry, even for K^+^.

It appears that the ion transfer mechanisms cannot be so easily classified as direct, or hard, or indirect or soft, but require consideration of the side chains of the amino acids that are thoroughly conserved. Water is present near the ions, but can be pushed somewhat to the side (albeit possibly not normally so, with K^+^ In WT). We have seen how the ion may be coordinated to multiple water molecules. Because of the role of the charges on the oxygens that solvate the ions, including K^+^ in WT, it may not be entirely fair to describe the mechanism as simply “knock-on”. KWK configurations are not all the same, as the water may sometimes be more strongly hydrogen bonded to the amino acid side chains than connected to the ion. A detailed examination of the orientation of water, charges on the atoms with consequent electrostatic forces, and symmetry, is required to understand rectification and selectivity in the SF. It would be interesting to calculate a Kir channel to see whether the electrostatic gradient is reversed, and whether or not the ratchet and pawl remain.

## CONCLUSIONS

1. The transfer of an ion from the cavity of the K_v_1.2 channel to the SF is a complex problem in re-solvation, involving transfer of hydrogen bonds and switching of electrostatic interactions. The co-solvation of the ion by threonines and water is necessary for the ion to move through the SF.
2. The problem is solved for K^+^ by the pore structure, with a consequent electrostatic gradient driving the ion forward, and the orientation of a water acting as the pawl in a ratchet and pawl that prevents a return of the ion (see (6) below).
3. Na^+^ goes slightly off the axis by enough to be more strongly bound electrostatically by the oxygens on one side of the pore than the other, as well as by tightly bound hydrating water molecules at the top of the cavity when the ion is in the S4 SF position. This makes it more difficult to move up, as there is a substantial barrier at the entrance to the SF.
4. It is a rather different problem for NH_4_^+^ which itself is not spherically symmetric. It is not as tightly held as Na^+^, but does not move as easily as K^+^, in part because the ion can form hydrogen bonds, unlike either Na^+^ or K^+^. It acts almost like a charged water.
5. The threonines are the right size to solvate the K^+^ on the pore axis; substituting serines leads to another form of asymmetry, as the ion can no longer be fully solvated on the pore axis.
6. The O(T373 –C=O) – O(T374 –OH) – H(T374 –OH) angle may be as important as electrostatics in making the “knock-on” mechanism possible, by acting as a pawl in a ratchet and pawl mechanism, allowing the knock on to be unidirectional, with no back flow (except for thermal fluctuations, which are necessary to comply with the Second Law of Thermodynamics (65)).
7. Taken together, these considerations suggest that the SF, and the pore as a whole, is a finely tuned structure. The requirement for fine tuning suggests in turn the reason for the conservation of the SF amino acid sequence through perhaps a billion years of evolutionary time. How much of this is related to C-type inactivation will require a study of the complete SF, which remains for future work; most probably, this too will require consideration of a combination of subtle effects.

## ACKNOWLEDGMENTS

This research used resources of the Center for Functional Nanomaterials, which is a U.S. DOE Office of Science Facility, and the Scientific Data and Computing Center, a component of the Computational Science Initiative, at Brookhaven National Laboratory under Contract No. DE-SC0012704, and resources at the High Performance Computation facility at City University of New York. Other funding was provided by the corresponding author.

## REFERENCES

1. Moroni A, Thiel G. Flip-flopping salt bridges gate an ion-channel. Nat Chem Biol. 2006;2:572–3.

2. Doyle DA, Cabral JM, Pfuetzner RA, Kuo A, Gulbis JM, Cohen SL, et al. The structure of the potassium channel: molecular basis of K^+^ conduction and selectivity. Science. 1998;280:69–77.

3. Jiang YL, A., Chen J, Cadene M, Chait BT, MacKinnon R. The open pore conformation of potassium channels. Nature. 2002;417:523–6.

4. Zhou Y, MacKinnon R. Ion binding affinity in the cavity of the KcsA potassium channel. Biochem. 2004;43:4978–82.

5. Zhou Y, Morais-Cabral JH, Kaufman A, MacKinnon R. Chemistry of ion coordination and hydration revealed by a K^+^ channel-FAB complex at 2.0 A resolution. Nature. 2001;414:43–8.

6. Kariev AM, Green ME. Quantum mechanical calculations on selectivity in the KcsA channel: the role of the aqueous cavity. J Phys Chem B. 2008;112:1293–8.

7. Kariev AM, Green ME. Quantum calculations on water in the KcsA channel cavity with permeant and non-permeant ions. Biochim Biophys Acta (Biomembranes). 2009;1788:1188–92.

8. Ban F, Kusalik P, Weaver DF. Density Functional Theory Investigations on the Chemical Basis of the Selectivity Filter in the K+ Channel Protein. J Am Chem Soc. 2004;126(14):4711–6.

9. Varma S, Rogers DM, Pratt LR, Rempe SB. Design principles for K+ selectivity in membrane transport. J Gen’l Physiol. 2011;137:479–88.

10. Eisenberg B. Interacting Ions in Biophysics: Real is not Ideal. Biophys J. 2013;104(9):1849–66.

11. Liu S, Lockless SW. Equilibrium selectivity alone does not create K+selective ion conduction in K+ channels. Nat Commun. 2013;4:3746/1-/7.

12. Ngo V, Stefanovski D, Haas S, Farley RA. Non-equilibrium dynamics contribute to ion selectivity in the KcsA channel. PLoS One. 2014;9(1):e86079.1-e/12, 12 pp.

13. Sauer DB, Zeng W, Raghunathan S, Jiang Y. Protein interactions central to stabilizing the K+ channel selectivity filter in a four-sited configuration for selective K+ permeation. Proc Natl Acad Sci U S A. 2011;108(40):16634–9, S/1-S/7.

14. Thomas M, Jayatilaka D, Corry B. An entropic mechanism of generating selective ion binding in macromolecules. PLoS Comput Biol. 2013;9(2):e1002914.

15. Wang S, Lee S-J, Maksaev G, Fang X, Zuo C, Nichols CG. Potassium channel selectivity filter dynamics revealed by single-molecule FRET. Nat Chem Biol. 2019:Ahead of Print.

16. Chaudhari MI, Rempe SB. Strontium and barium in aqueous solution and a potassium channel binding site. J Chem Phys. 2018;148(22):222831/1-/7.

17. Eichmann C, Frey L, Maslennikov I, Riek R. Probing Ion Binding in the Selectivity Filter of the KcsA Potassium Channel. J Am Chem Soc. 2019;141(18):7391–8.

18. Langan PS, Vandavasi VG, Weiss KL, Coates L, Langan PS, Vandavasi VG, et al. Anomalous X-ray diffraction studies of ion transport in K(+) channels. Nat Commun. 2018;9(1):4540.

19. Hoomann T, Jahnke N, Horner A, Keller S, Pohl P. Filter gate closure inhibits ion but not water transport through potassium channels. Proc Natl Acad Sci U S A. 2013;110(26):10842–7,S/1-S/2.

20. Imai S, Osawa M, Takeuchi K, Shimada I. Structural basis underlying the dual gate properties of KcsA. Proc Natl, Acad Sci. 2010;107:6216–21.

21. Langan PS, Vandavasi VG, Sullivan B, Weiss K, Coates L, Harp J. Crystallization of a potassium ion channel and X-ray and neutron data collection. Acta Crystallogr F Struct Biol Commun. 2019;75(Pt 6):435–8.

22. Tilegenova C, Cortes DM, Jahovic N, Hardy E, Hariharan P, Guan L, et al. Structure, function, and ion-binding properties of a K+ channel stabilized in the 2,4-ion-bound configuration. Proc Natl Acad Sci U S A. 2019;116(34):16829–34.

23. Kratochvil HT, Carr JK, Matulef K, Annen AW, Li H, Maj M, et al. Instantaneous ion configurations in the K+ ion channel selectivity filter revealed by 2D IR spectroscopy. Science (Washington, DC, U S). 2016;353(6303):1040–4.

24. Strong SE, Hestand NJ, Kananenka AA, Zanni MT, Skinner JL. IR Spectroscopy Can Reveal the Mechanism of KD Transport in Ion Channels. Biophys J. 2020;118:254–61.

25. Weingarth M, van der Cruijsen EAW, Ostmeyer J, Lievestro S, Roux B, Baldus M. Quantitative analysis of the water occupancy around the selectivity filter of a K+ channel in different gating modes. J Am Chem Soc. 2014;136(5):2000–7.

26. Cifuentes AA, Semiao FL. Quantum model for a periodically driven selectivity filter in a K+ ion channel. J Phys B: At, Mol Opt Phys. 2014;47(22):225503/1-/6, 6 pp.

27. Summhammer J, Sulyok G, Bernroider G. Quantum dynamics and non-local effects behind ion transition states during permeation in membrane channel proteins. Entropy. 2018;20(8):558/1-/13.

28. Vaziri A, Plenio MB. Quantum coherence in ion channels: resonances, transport and verification. arXivorg, e-Print Arch, Quantum Phys. 2010:1–10, 1006.3892v1 [quant-ph].

29. De S, March N, Prado SD, Brunnet LG. Coulomb interaction rules timescales in potassium ion channel tunneling. J Phys: Condens Matter. 2018;30(25):255101/1-/6.

30. De S, Rinsha CH, Thamleena A H, Joseph A, Ben A, V. U K. Roles of different amino-acid residues towards binding and selective transport of K+ through KcsA K+-ion channel. Phys Chem Chem Phys. 2018;20(25):17517–29.

31. Pichierri F. Macrodipoles of potassium and chloride ion channels as revealed by electronic structure calculations. J Mol Struct: THEOCHEM. 2010;950(1-3):79–82.

32. Illingworth CJ, Domene C. Many-body effects and simulations of potassium channels. Proc R Soc A. 2009;465(2106):1701–16.

33. Illingworth CJR, Furini S, Domene C. Computational Studies on Polarization Effects and Selectivity in K+ Channels J Chem Theory and Computation. 2010;6:3780–92.

34. Pless SA, Galpin JD, Niciforovic AP, Kurata HT, Ahern CA. Hydrogen bonds as molecular timers for slow inactivation in voltage-gated potassium channels. eLife. 2013;2:e01289.1-e/14.

35. Miloshevsky GV, Jordan PC. Conformational changes in the selectivity filter of the open-state KcsA channel: An energy minimization study. Biophys J. 2008;95:3239–51.

36. Bucher D, Guidoni L, Carloni P, Rothlisberger U. Coordination numbers of K(+) and Na(+) Ions inside the selectivity filter of the KcsA potassium channel: insights from first principles molecular dynamics Biophys J. 2010;98:L47–L9.

37. Cifuentes AA, Semiao FL. Quantum model for the selectivity filter in K+ ion channel. arXivorg, e-Print Arch, Phys. 2013:1–4, 1312.4056v1 [physics.bio-ph].

38. Das B, Banerjee K, Gangopadhyay G. Entropy hysteresis and nonequilibrium thermodynamic efficiency of ion conduction in a voltage-gated potassium ion channel. Phys Rev E: Stat, Nonlinear, Soft Matter Phys. 2012;86(Copyright (C) 2013 American Chemical Society (ACS). All Rights Reserved.):061915/1-/10.

39. Wawrzkiewicz-Jalowiecka A, Dworakowska B, Grzywna ZJ. The temperature dependence of the BK channel activity - kinetics, thermodynamics, and long-range correlations. Biochim Biophys Acta, Biomembr. 2017;1859(10):1805–14.

40. Wawrzkiewicz-Jalowiecka A, Grzywna ZJ. The role of entropic potential in voltage activation and K+ transport through Kv1.2 channels. J Chem Phys. 2018;148(11):115103/1-/10.

41. Portella G, Hub J, Vesper M, deGroot B. Not only enthalpy: Large Entropy Contribution to Ion Permeation Barriers in Single File Channles. Biophys J 2008;95:2275–82.

42. Roux B, Berneche S, Egwolf B, Lev B, Noskov SY, Rowley CN, et al. Ion selectivity in channels and transporters. J Gen Physiol. 2011;137(5):415–26.

43. Dixit PD, Asthagiri D. Thermodynamics of ion selectivity in the KcsA K+ channel. J Gen Physiol. 2011;137(5):427–33.

44. Kraszewski S, Boiteux C, Ramseyer C, Girardet C. Determination of the charge profile in the KcsA selectivity filter using ab initio calculations and molecular dynamics simulations. Phys Chem Chem Phys. 2009;11:8606–13.

45. Shrivastava IH, Tieleman DP, Biggin PC, Sansom MSP. K+ versus Na+ ions in a K channel selectivity filter: A simulation study. Biophys J. 2002;83(2):633–45.

46. Yu H, Roux B. On the utilization of energy minimization to the study of ion selectivity. Biophys J. 2009;97(8):L15–L7.

47. Grottesi A, Domene C, Haider S, Sansom MSP. Molecular dynamics simulation approaches to K channels: conformational flexibility and physiological function. IEEE Trans Nanobioscience. 2005;4(1):112–20.

48. Shrivastava IH, Capener CE, Forrest LR, Sansom MSP. Structure and dynamics of K channel pore-lining helices: a comparative simulation study. Biophys J. 2000;78(Copyright (C) 2012 American Chemical Society (ACS). All Rights Reserved.):79–92.

49. Alam A, Jiang Y. Structural analysis of ion selectivity in the NaK channel. Nat Struct Mol Biol. 2009;16(1):35–41.

50. Alam A, Jiang Y. High-resolution structure of the open NaK channel Nature Struct Molec Biol. 2009;16:30–4.

51. Brettmann JB, Urusova D, Tonelli M, Silva JR, Henzler-Wildman KA. Role of protein dynamics in ion selectivity and allosteric coupling in the NaK channel. Proc Natl Acad Sci U S A. 2015;112(50):15366–71.

52. Derebe MG, Sauer DB, Zeng W, Alam A, Shi N, Jiang Y. Tuning the ion selectivity of tetrameric cation channels by changing the number of ion binding sites. Proceedings National Academy Sci, US. 2011;108:598–602.

53. Shen R, Guo W. Mechanism for Variable Selectivity and Conductance in Mutated NaK Channels. J Phys Chem Lett. 2012;3(19):2887–91.

54. Rossi M, Tkatchenko A, Rempe SB, Varma S. Role of methyl-induced polarization in ion binding. Proc Natl Acad Sci U S A. 2013;110(32):12978–83.

55. Kariev AM, Green ME. Quantum Calculation of Proton and Other Charge Transfer Steps in Voltage Sensing in the Kv1.2 Channel. J Phys Chem B. 2019;123:7984–98.

56. Oakes V, Furini S, Domene C. Insights into the Mechanisms of K+ Permeation in K+ Channels from Computer Simulations. J Chem Theory Comput. 2019.

57. Kariev AM, Znamenskiy VS, Green ME. Quantum mechanical calculations of charge effects on gating the KcsA channel. Biochem Biophys Acta(Biomembranes). 2007;1768:1218–29.

58. Nimigean C, Miller C. Na^+^ block and permeation in a K^+^channel of known structure. J Gen’l Physiol. 2002;120:323–35.

59. Hille B. Potassium Channels in Myelinatedd Nerve. J Gen’l Physiol. 1973;61:669–86.

60. Frisch MJ, Trucks GW, Schlegel HB, Scuseria GE, Robb MA, Cheeseman JR, et al. Gaussian 09, Revision D.01. Wallingford CT: Gaussian, Inc.; 2009.

61. Weinhold F. NBO 5.0 Program Manual. Madison, WI: Theoretical Chemistry Institute, U. of Wisconsin; 2001.

62. Weinhold F. Natural Bond Orbital Analysis: A Critical Overview of Relationships to Alternative Bonding Perspectives. J Comp Chem. 2012;33:2363–79.

63. Chen X, Wang Q, Ni F, Ma J. Structure of the full-length Shaker potassium channel Kv1.2 by normal-mode-based X-ray crystallographic refinement. Proc Natl Acad Sci. 2010;107:11352–7.

64. Thompson AN, Kim I, Panosian TD, Iverson TM, Allen TW, Nimigean CM. Mechanism of potassium-channel selectivity revealed by Na(+) and Li(+) binding sites within the KcsA pore Nature Struct Molec Biol. 2009;16:1317–24.

65. Feynman RP, Leighton RB, Sands M. Feynman Lectures on Physics. Boston: Addison-Wesley; 1964.

